# INO80 rapidly shuttles nucleosomes between chromatin barriers

**DOI:** 10.64898/2026.09.20.752975

**Authors:** Benjamin Ambrose, Paul Girvan, Michael T. Skehan, Elizabeth A. McCormack, Deborah I. Egharevba, David S. Rueda

## Abstract

ATP-dependent chromatin remodelers establish nucleosome organization, but how they move nucleosomes over extended distances and respond to neighboring chromatin remains unclear. Here, we combine correlative optical tweezers and fluorescence microscopy, and single-molecule FRET to visualize human INO80 activity on naked and chromatinized DNA. hINO80 undergoes free one-dimensional diffusion along DNA with multiple, nucleotide-regulated, diffusive states. Upon engaging a nucleosome, hINO80 drives rapid, processive nucleosome sliding over thousands of base pairs in bidirectional bursts. Neighboring nucleosomes prevent passage and redirect translocation, confining mobile nucleosomes to repeated movement between chromatin boundaries. Single-molecule FRET reveals transitions consistent with INO80 switching between opposing nucleosome-binding orientations, while fluorescence stoichiometry and mass photometry show that two hINO80 complexes can simultaneously occupy one nucleosome. These findings identify non-exclusive mechanisms for directional reversal and reveal how long-range hINO80 translocation is converted into boundary-constrained nucleosome repositioning, providing a dynamic framework for understanding nucleosome organization within chromatin.

---

From yeast to humans, transcription, replication, DNA damage detection and repair require access to nuclear DNA. This access is controlled by the positioning, composition and higher-order organization of nucleosomes within chromatin (Luger *et al*., 1997; Kornberg *et al*., 2020). ATP-dependent chromatin remodellers regulate this by sliding or evicting nucleosomes and altering their histone composition^1^. Their catalytic subunits contain an ATPase of the Snf2 family^2^ and are classified, according to their surrounding accessory domains, into the ISWI, CHD, SWI/SNF and INO80 families^3^. The INO80 family is distinctive because it includes members that catalyze either nucleosome sliding (INO80), or histone-variant exchange (SWR1/SRCAP)^4–6^.

INO80 is a large (>1 MDa), multi-subunit complex that slides and spaces nucleosomes *in vitro*^4,7,8^ and generates longer inter-nucleosomal spacing than other remodellers^9^. In cells, INO80 localizes to transcription start sites, where it associates with the +1 nucleosome^10^. The +1 nucleosome defines the downstream phasing of gene-body nucleosomes and lies adjacent to an upstream nucleosome-depleted region (NDR) of approximately 100-1000 bp^11,12^. INO80 has been implicated in establishing and maintaining this organization, but how its nucleosome-sliding activity generates such patterns remains unclear. INO80 also participates in several DNA-repair pathways, where nucleosomes must be repositioned to expose damaged DNA and facilitate access by repair factors^13–15^. However, the underlying remodelling mechanisms remain poorly understood.

Cryo-electron microscopy has revealed the architecture of INO80 bound to a nucleosome^9,16,17^. A RUVBL1/RUVBL2 heterohexamer forms a scaffold around which the catalytic subunit (INO80) and associated co-factors are organized (**Fig. 1A**). The INO80 motor binds nucleosomal DNA near the entry site, at superhelical location 6/7 (SHL6/7), while ARP5/IES6 engages the opposite side of the nucleosome near SHL2/3 and couples ATP hydrolysis to DNA translocation^18–20^. This allows INO80 to “push” entry DNA into the nucleosome, in contrast with most Snf2-family remodellers, including SWR1, which bind at SHL2 and “pull” DNA around the histone core. INO80 can also slide hexasomes, although this mechanism remains to be elucidated^20–23^.

**Figure 1.**
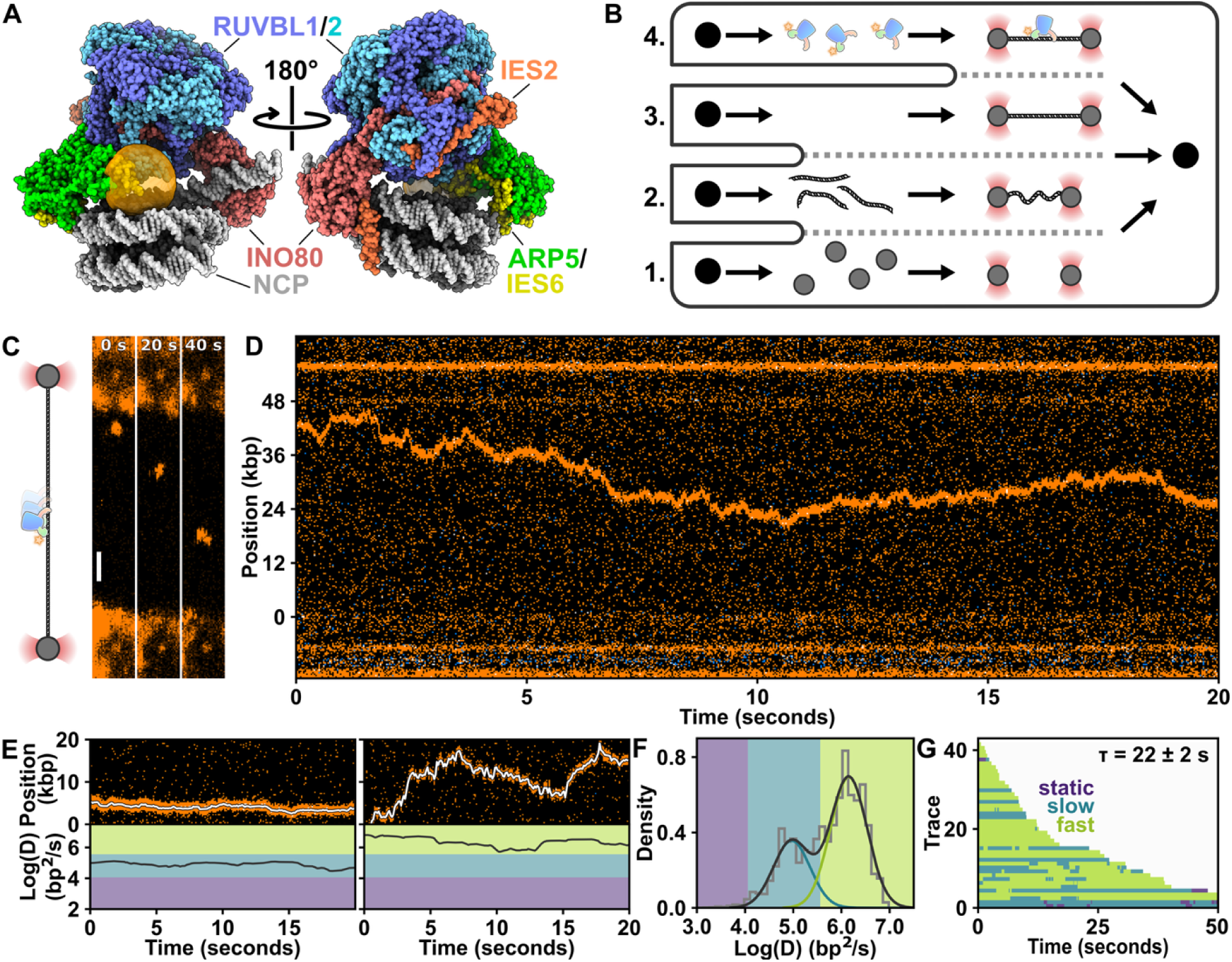
hINO80 diffuses rapidly on naked DNA. **A.** Structure of hINO80 on a nucleosome, with location of Atto 647N label (orange cloud). **B.** Optical tweezers laminar flow cell with streptavidin beads (Channel 1), biotinylated λ-DNA (Channel 2), imaging buffer (Channel 3) and labelled hINO80 (Channel 4). **C.** Schematic of hINO80 on optically trapped DNA, and 2D frames (**Supplementary Movie 1**) of hINO80 diffusing (scale bar 1 μm). **D.** Kymograph of hINO80 diffusing. **E.** Example kymographs of fast and slow diffusion, with a rolling diffusion analysis below (fast (green), slow (blue), and static (purple) regimes). **F.** Rolling diffusion coefficient histogram (grey) with log-Gaussian fits (black) (N = 46). **G.** Rastergram of traces (N = 46) shaded according to their diffusion regime as **E**. Mean lifetime on the DNA is 22 ± 2 s (standard deviation, SD).

Using isolated nucleosomes positioned near the ends of short DNA fragments, previous single-molecule studies have shown that INO80 processively moves nucleosomes towards the DNA centre. Translocation occurs in rapid 8-12 bp bursts, with a rate that depends on the length of the flanking DNA^24^. These experiments established important features of short-range nucleosome sliding but could not reveal how INO80 moves nucleosomes over much greater distances. In particular, how INO80 behaves when a translocating nucleosome encounters neighbouring nucleosomes, as it would at the boundaries of an NDR, remains unknown.

To address this question, we have combined optical tweezers and single-molecule fluorescence microscopy experiments^25^ to investigate how the human INO80 complex (hINO80) behaves on chromatinized DNA. The data show that hINO80 can rapidly translocate nucleosomes over thousands of base pairs, substantially farther and faster than previously observed. Furthermore, hINO80 cannot translocate a nucleosome past other static nucleosomes on the DNA. Instead, static nucleosomes act as translocation barriers at which hINO80 stops and reverses direction. When confined between two static nucleosomes, hINO80 slides the intervening nucleosome repeatedly back and forth between the two static barrier nucleosomes.

Single-molecule FRET measurements reveal that hINO80 can switch between opposite sides of a nucleosome, similar to its relative SWR1^26^, providing a mechanism for directional reversal. These data also show that two hINO80 complexes can simultaneously occupy a single nucleosome, suggesting an additional mechanism whereby oppositely oriented complexes could drive movement in opposite directions. Together, our findings establish how hINO80 carries out long-range nucleosome sliding within chromatin, and suggest a mechanism by which it can slide nucleosomes between fixed boundaries, including those surrounding transcription start sites.

## Results

### hINO80 diffuses freely on naked DNA

First, we sought to characterize the one-dimensional (1D) diffusion of hINO80 along naked DNA using correlative optical tweezers and fluorescence microscopy^25^. Such diffusion likely facilitates the search for nucleosomal substrates across nucleosome-depleted regions. We purified the conserved core human INO80 complex (hINO80)^19^ lacking ARP5/IES6 (ΔARP5/IES6), and reconstituted it with an ARP5/IES6 module fluorescently labelled via a ybbR tag conjugated Atto 647N^27^ (**Fig. 1A** and **Supplementary Fig. 1A**). Reconstitution with labelled ARP5/IES6 did not measurably alter the nucleosome-sliding activity of the complex (**Supplementary Fig. 1B**).

Biotinylated λ-DNA was tethered between two optically trapped streptavidin-coated beads, briefly incubated in channel containing labelled hINO80 and transferred to a protein-free imaging channel (**Fig. 1B**). hINO80 trajectories were recorded either by two-dimensional imaging (**Fig. 1C** and **Supplementary Movie 1)** or as one-dimensional kymographs (**Fig. 1D**). Approximately two-thirds of the time, the examined complexes diffused rapidly over long distances, with some traversing approximately half of the λ-DNA molecule (∼24 kbp) within 10 s (**Fig. 1D**). The remaining third displayed markedly slower diffusion (**Fig. 1E**).

To quantify this heterogeneity, we calculated local diffusion coefficients using a rolling-window analysis^28,29^. The resulting distribution was bimodal, revealing fast (∼10^6^ bp^2^/s) and slow (∼10^5^ bp^2^/s) diffusive states (**Fig. 1E, F**). A rastergram of the local diffusion coefficients showed that most complexes predominantly occupied the fast state, although a subset (N = 12/46) switched between the two states within individual trajectories (**Fig. 1G**). Because hINO80 contains multiple DNA-binding elements^16,17,30,31^, the two states may reflect different degrees or configurations of DNA engagement.

Under standard conditions, hINO80 remained bound to λ-DNA for an average of 22 ± 2 s before dissociating (**Fig. 1G**). Decreasing the KCl concentration to 25 mM extended the lifetime to 30 ± 4 s, and increased the proportion of slow-diffusing complexes (**Supplementary Fig. 2A**). Conversely, increasing the KCl concentration to 175 mM shortened this lifetime to 6 ± 1 s and increased the proportion of fast-diffusing complexes (**Supplementary Fig. 2C**). Thus, electrostatic interactions stabilize hINO80 binding to DNA and favour the slow-diffusing state (**Supplementary Fig. 2D** and **E**), consistent with more extensive engagement of the complex with the DNA.

Next, we examined how nucleotide state affects hINO80 diffusion. ATP increased the mean DNA-bound lifetime to 28 ± 4 s and converted the bimodal distribution into an apparently single population with an intermediate diffusion coefficient (**Supplementary Fig. 3A**). The non-hydrolyzable ADP·BeF□, which mimics an ATP-bound transition state, further extended the lifetime to 42 ± 5 s and produced a distinct population of immobile, DNA-bound complexes that was not detected under any other condition (**Supplementary Fig. 3B**). By contrast, ADP reduced the lifetime to 11 ± 2 s and shifted both diffusive populations towards higher diffusion coefficients (**Supplementary Fig. 3C**). Finally, mean-squared-displacement analysis yielded anomalous diffusion exponents (α) centered near 1 both in the presence and absence of ATP (**Supplementary Fig. 3D**), consistent with free diffusion rather than directed motion.

Together, these results show that, on naked DNA, hINO80 undergoes free 1D diffusion and dynamically occupies at least two diffusive states. Salt and nucleotide conditions regulate both the mobility and DNA-bound lifetime of the complex, suggesting that its diffusive behavior is controlled by changes in the extent or configuration of DNA engagement.

### hINO80 slides nucleosomes rapidly, processively and bidirectionally

To examine hINO80 activity on chromatinized DNA, we assembled human nucleosomes directly on tethered λ-DNA using purified histone octamers containing fluorescently labelled H2A (**Supplementary Fig. 4A**) and yeast Nap1 (yNAP1) as an assembly chaperone^32^. DNA tethers held at low force (∼1 pN) were incubated with histone octamers and yNap1 in a separate channel of the laminar-flow cell (**Supplementary Fig. 4B**). At this force, the DNA linkers exiting the nucleosome are expected to cross and emerge in opposite directions, maintaining their native DNA-wrapped structure^33^. Histone concentration and incubation time were optimized to generate approximately 3-10 nucleosomes per tether. This approach produced nucleosomes at stochastic positions, thereby enabling investigation of INO80 activity across a range of nucleosomal contexts.

We first validated the assembled nucleosomes using force spectroscopy. Increasing the tension on the DNA produced discrete rupture events accompanied by increases in DNA extension (**Fig. 2A**), as expected for nucleosome unwrapping^34^. Fitting the force-extension curves before and after rupture with an extensible worm-like chain model yielded a mean contour-length increase of ∼24 nm per nucleosome, consistent with previous measurements^34–36^. Across 91 force-extension traces, nucleosome unwrapping occurred between 10 and 35 pN and produced the expected contour-length increments (**Fig. 2B**). Most traces contained a single unwrapping event (N = 79), whereas 9, 2 and 1 traces contained two, three and four coincident events, respectively.

**Fig 2.**
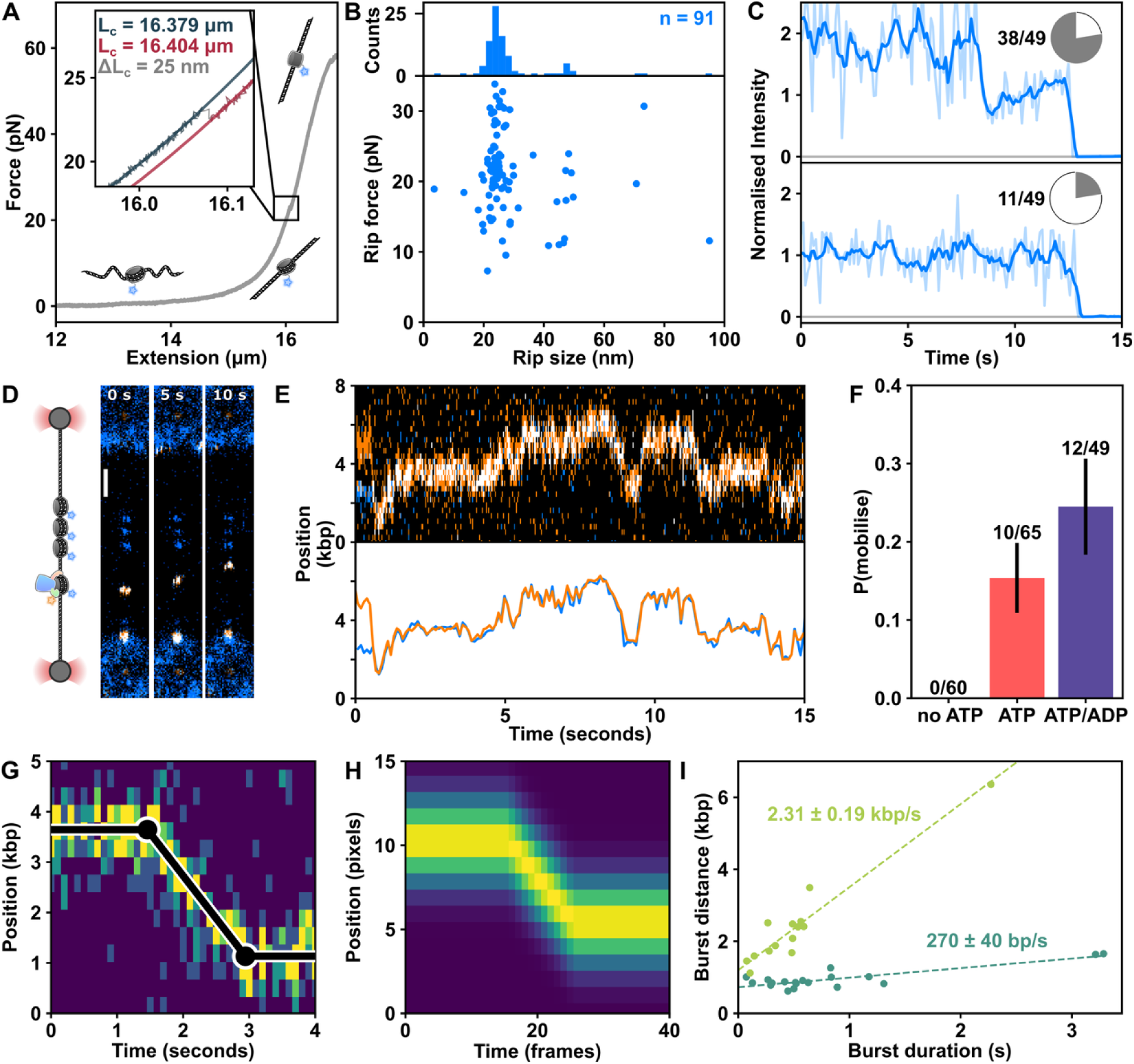
hINO80 slides nucleosomes rapidly, processively and bidirectionally. **A.** Force extension curve of a nucleosome on λ-DNA. Inset: change in contour length (ΔLc) upon unwrapping. **B.** Nucleosome unwrapping force as a function of ΔLc (n = 91), showing expected ∼24 nm contour length increase. **C.** Fluorescence trajectories of H2A-AF555 nucleosome showing 78% (N = 38/49) two-step photobleaching. **D.** Schematic of hINO80 (orange) and nucleosomes (blue) on λ-DNA with 3 frames showing nucleosome sliding (white) (**Supplementary Movie 2**). **E.** Kymograph (top) and localization (bottom) of hINO80 (red) diffusing into a nucleosome (blue) and subsequent bidirectional sliding. **F.** Fraction of sliding nucleosomes in the absence and presence of ATP or ATP with ADP (error bars are binomial SD). **G.** Nucleosome sliding burst (∼2.5 kbp in ∼1.5 s) fit with piecewise function (black). **H.** Idealized piecewise function used to fit bursts. **I.** Burst distance as a function of burst time reveals two speeds of hINO80-mediated nucleosome sliding.

Photobleaching analysis further confirmed that the assembled particles predominantly contained complete histone octamers (**Fig. 2C**). Of 49 nucleosomes analysed, 38 (78%) exhibited two photobleaching steps, as expected for particles containing two labelled H2A-H2B dimers. The remaining 11 particles (22%) exhibited a single step. This distribution is consistent with the estimated H2A labelling efficiency of approximately 90%, which predicts that almost all the nucleosomes should contain two labelled H2A molecules. A substantial population of hexasomes or tetrasomes would instead have produced a considerably lower proportion of two-step trajectories.

We next introduced fluorescently labelled hINO80 to the chromatinized DNA in the presence of ATP (**Fig. 2D** and **Supplementary Movie 2**). hINO80 colocalized with individual nucleosomes and translocated with them rapidly, processively and bidirectionally over several thousand base pairs within seconds (**Fig. 2E**). Adding ADP at a physiological ATP ratio (1 mM ATP and 0.1 mM ADP) significantly increased the proportion of nucleosome-bound hINO80 complexes that underwent translocation (**Fig. 2F**). No hINO80-nucleosome translocation was detected in the absence of ATP (**Fig. 2F**). Under this condition, hINO80 diffused along the DNA and bound nucleosomes but neither displaced nor bypassed them (**Supplementary Fig. 4C**). By contrast, ATP occasionally enabled hINO80 to bypass a nucleosome without translocating it (**Supplementary Fig. 4D** and **E**). These observations indicate that ATP hydrolysis is required both for nucleosome translocation and for INO80 to overcome the nucleosome barrier encountered during diffusion.

Nucleosome translocation occurred as rapid, bidirectional bursts separated by stationary pauses (**Fig. 2G**). To quantify this behavior, we fitted a piecewise stationary-linear-stationary model, incorporating a one-dimensional Gaussian point-spread function, to 37 clearly resolved translocation bursts from the ATP and ATP-ADP datasets (**Fig. 2H**). The fit identified the transitions between stationary and mobile states, allowing the start and end of each burst to be localized in both space and time. From these changepoints, we calculated the burst distance and burst duration of each translocation event (mean distance 1.0 ± 0.2 kbp, mean duration 0.6 ± 0.1 s (**Supplementary Fig. 4F**)). To quantify the sliding rate of these events, we plotted burst distance as a function of burst duration, which revealed two populations of hINO80-mediated nucleosome sliding, with speeds of 270 ± 40 bp/s and 2.3 ± 0.2 kbp/s (**Fig. 2I**). Thus, on extended chromatinized substrates, hINO80 slides nucleosomes substantially farther and faster than previously observed using short DNA substrates^4,7,19,24^.

Together, these results establish that hINO80 is a highly processive nucleosome motor that translocates nucleosomes over kilobase distances in rapid, ATP-dependent bursts and can reverse direction between successive bursts.

### Flanking nucleosomes constrain and reverse hINO80 translocation

In a subset of trajectories, an hINO80-translocated nucleosome was confined between two static nucleosomes separated by several kilobase pairs. The mobile hINO80-nucleosome complex repeatedly shuttled back and forth between these flanking nucleosomes, which acted as physical barriers to further translocation (**Fig. 3A** and **Supplementary Fig. 4G**). The complex also paused at intermediate locations, often returning to the same positions within an individual trajectory. However, it never bypassed either boundary nucleosome. Thus, neighboring nucleosomes can delimit the range of INO80-mediated translocation and promote repeated directional switching.

**Figure 3.**
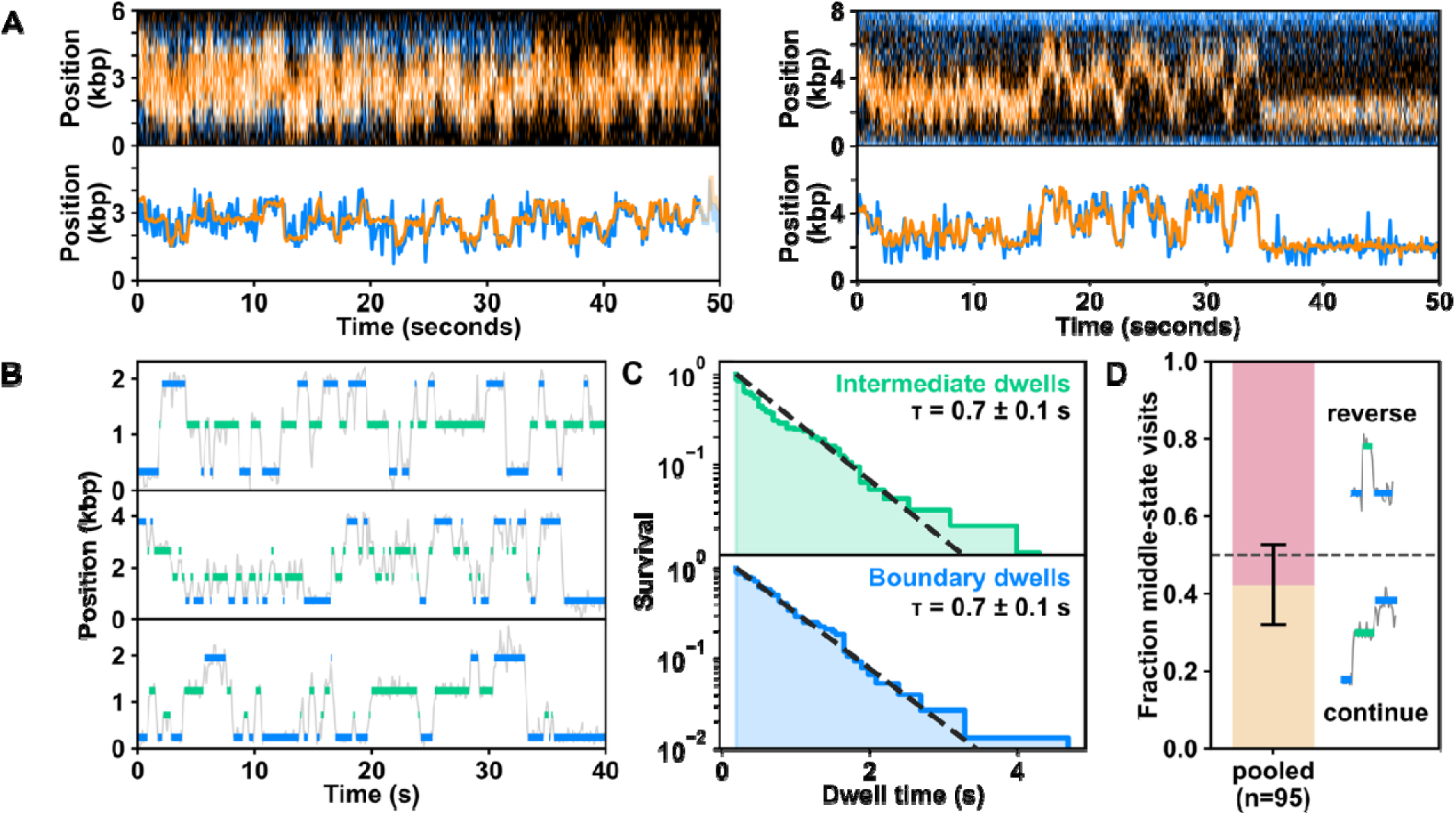
Flanking nucleosomes constrain and reverse hINO80 translocation. **A.** Kymographs (top) and localizations (bottom) of hINO80 (orange) repeatedly sliding a nucleosome back and forth between two flanking nucleosomes (blue), exhibiting regular pauses at intermediate locations. **B.** K-means clustering analysis of 3 kymographs with shuttling nucleosomes between boundaries (blue) and intermediate (green) pausing locations. **D.** Dwell time distributions of intermediate (top) and boundary positions (bottom), and single exponential fits (dashed). **E.** Bar chart showing hINO80-nucleosome complex direction following intermediate pauses resulting in either reversal (pink) or continuation (yellow) events.

To quantify this behavior, we used k-means clustering to identify the positions of the boundary nucleosomes and recurrent intermediate pauses (**Fig. 3B** and **Methods**). Each trajectory contained only one or two recurrent intermediate pausing positions. Dwell-time distributions at both intermediate and boundary positions followed single-exponential kinetics, with identical half-lives of 0.7 ± 0.1 s (**Fig. 3C**). This similarity suggests that departure from both types of pauses is governed by the same rate-limiting step.

Following an intermediate pause, the hINO80-nucleosome complex either resumed movement in its original direction or reversed direction. Of 95 intermediate pauses, 40 were followed by continued forward movement and 55 by reversal (**Fig. 3D**). This difference was not statistically significant (two-sided binomial test, *p = 0.151*), indicating that direction after an intermediate pause is effectively unbiased. Directional reversal is, therefore, an intrinsic feature of hINO80 translocation and does not require collision with a boundary nucleosome, although such collisions necessarily constrain subsequent movement to the opposite direction.

Previous single-molecule FRET studies have shown that hINO80 can pause and reverse while moving an isolated nucleosome along a short DNA substrate^24,37^. Our observations extend this behavior to a chromatinized substrate and show that hINO80 can repeatedly shuttle a nucleosome between neighboring nucleosomal barriers over kilobase distances. Rather than acting solely as passive steric barriers, neighboring nucleosomes may define the effective ends of the available linker DNA, restricting the region over which the intervening nucleosome can move. Such confined, bidirectional translocation provides a potential mechanism for nucleosome spacing: as neighboring nucleosomes become more closely positioned in nuclear chromatin, repeated movement and reversal between these boundaries could progressively redistribute the intervening nucleosome towards an average central position.

Together, these results provide a dynamic view of how hINO80 activity is constrained by chromatin architecture and suggest that barrier-induced confinement and local flanking-DNA sensing cooperate to organize nucleosome spacing.

### INO80 dynamics and dual occupancy suggest mechanisms for directional reversal

The ability of hINO80 to reverse nucleosome translocation without detectable dissociation raises the interesting question of how the complex changes direction. We considered two non-exclusive mechanisms. First, a single hINO80 complex could switch between opposing binding orientations on the nucleosome. We previously showed that the related SWR1 remodeler switches between opposite sides of a nucleosome, contributing to its processivity and substrate specificity^26^. Other remodelers, including Chd1 and human BAF, can also engage nucleosomes in opposing orientations associated with bidirectional sliding^38,39^.

To test whether hINO80 undergoes similar dynamics, we immobilized mono-nucleosomes containing an internal Alexa 555 donor fluorophore at SHL-2.5 and incubated them with Atto 647N-labelled INO80 as the FRET acceptor, in the absence of nucleotide (**Fig. 4A**). Alternating-laser excitation (ALEX) allowed INO80 binding and the resulting FRET dynamics to be monitored simultaneously^26,40^. The donor position was selected to generate distinguishable FRET efficiencies when hINO80 occupied opposing nucleosome-binding orientations.

**Figure 4.**
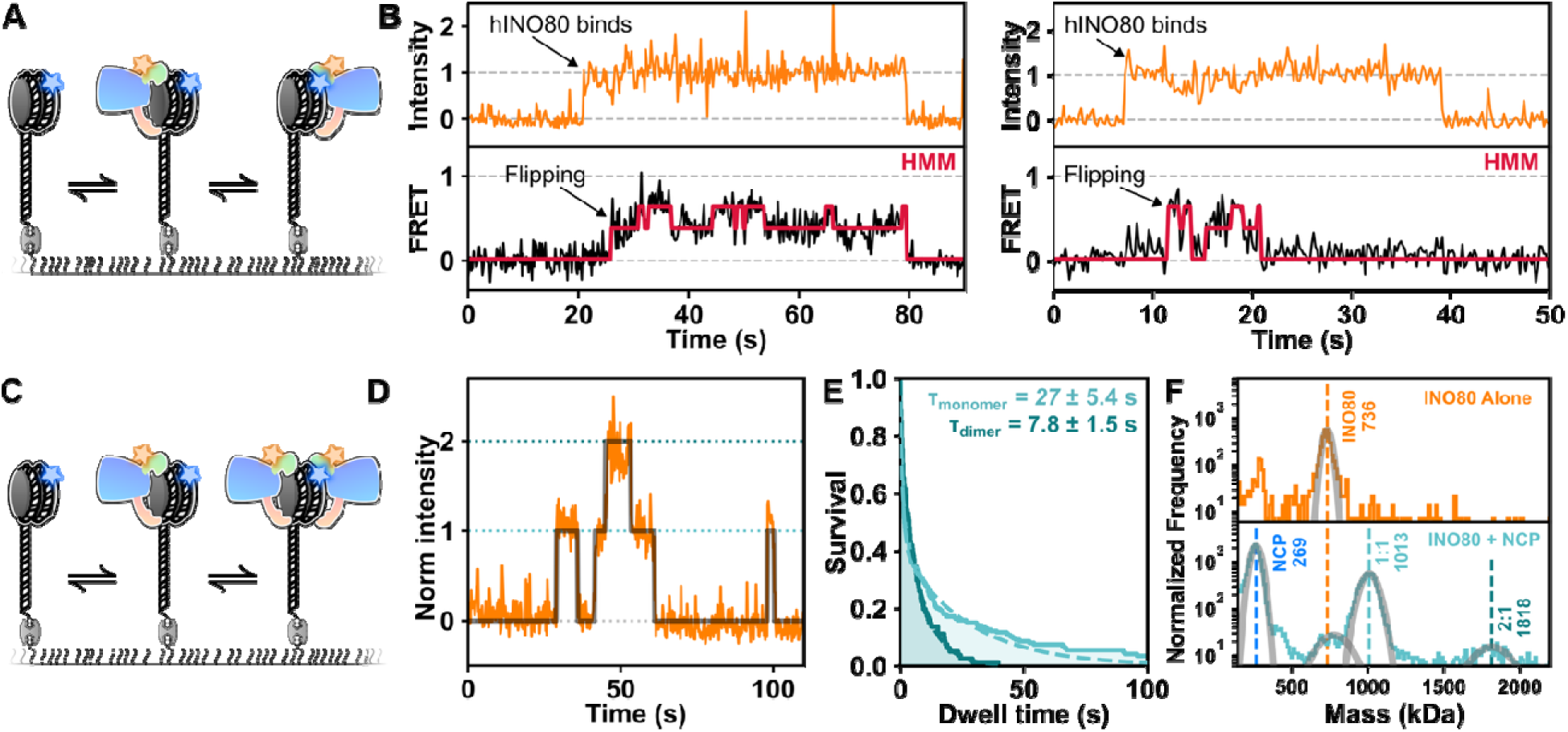
INO80 dynamics and dual occupancy suggest mechanisms for directional reversal. **A.** Diagram of the smFRET experiment for nucleosome flipping. Surface-immobilized labelled nucleosomes can bind labeled hINO80 in two possible orientations distinguishable by FRET. **B.** Single-molecule trajectories showing initial hINO80 event (top) and subsequent FRET dynamics (bottom). **C.** Diagram of fluorescence intensity experiment to determine hINO80 stoichiometry. **D.** Single-molecule fluorescence trajectory of labeled hINO80 binding a surface-immobilized mono-nucleosome, showing sequential binding of two hINO80 (∼40 s). **E.** Dwell time distributions of the 1:1 (light green) and 2:1 (green) complexes with double exponential fits (dashed lines). **F.** Mass photometry spectra of hINO80 in the absence (top) and presence of nucleosomes (bottom), indicating the presence of 2:1 complex at 1818 kDa. Expected masses are: 278 kDa (nucleosome), 743 kDa (hINO80), 1021 kDa (1:1 complex) and 1764 kDa (2:1 complex).

A subset of trajectories (10 of 67) showed the arrival of a single INO80 complex, initially without detectable FRET, followed several seconds later by the appearance of FRET and subsequent rapid transitions between distinct FRET levels (**Fig. 4B** and **C** and **Supplementary Fig. 5A**). Hidden Markov modelling identified two predominant non-zero FRET states. Because the nucleosomal substrate contained a 113-bp DNA overhang, the initial zero-FRET state likely represents INO80 bound to the distal extranucleosomal DNA. The subsequent transitions between FRET states are consistent with hINO80 switching between alternative orientations on the nucleosome, although conformational rearrangements within a single binding orientation cannot be excluded. These dynamics provide a potential mechanism by which a nucleosome-bound INO80 complex could reverse the direction of translocation without dissociating from the substrate.

A second mechanism could involve two oppositely oriented hINO80 complexes bound simultaneously to the same nucleosome. Each complex would be positioned to drive translocation in a different direction, with directional switching resulting from a change in which motor is productively engaged. Consistent with this possibility, previous biochemical measurements showed that hINO80 remodels nucleosomes most efficiently at a ratio of 2:1^16^. Such an arrangement could also allow rapid reversal when a translocating nucleosome encounters a neighbouring nucleosomal barrier (**Fig. 3**).

To test whether two hINO80 complexes can occupy a single nucleosome, we analyzed the fluorescence intensity of labelled hINO80 bound to surface-immobilized mono-nucleosomes (**Fig. 4D**). In 26 of 67 trajectories, hINO80 fluorescence increased and decreased in discrete steps consistent with sequential binding of two complexes to one nucleosome (**Fig. 4E** and **Supplementary Fig. 5B** and **C**). In nearly all cases, the second intensity increase occurred after a detectable delay, indicating that the 2:1 complex formed through two independent binding events rather than association of a preformed INO80 dimer.

We used hidden Markov modelling to assign states containing zero, one or two INO80 complexes and determine their dwell-time distributions (**Fig. 4F** and **Supplementary Fig. 5D** and **E**). The distributions were biexponential, with a fast component (*τ*_fast_ < 1 s) that likely reflects transient interactions with the extranucleosomal overhang DNA. The slower component had a lifetime of 27 ± 5 s for the 1:1 INO80-nucleosome complex and 8 ± 2 s for the 2:1 complex. The shorter lifetime of the 2:1 state indicates that the second hINO80 complex is less stably bound. One possible explanation is that only one of the two complexes can form optimal contacts, including the A-module, with the long DNA overhang, although steric or conformational constraints within the 2:1 complex could also contribute.

Finally, we used mass photometry to determine whether 2:1 complexes also formed in solution, independently of surface immobilization (**Fig. 4G**). hINO80 alone produced a single peak at approximately 740 kDa, consistent with the mass of one hINO80 complex. Addition of nucleosomes produced three additional peaks at approximately 270, 1,010 and 1,820 kDa, corresponding to free mono-nucleosomes, 1:1 hINO80-nucleosome complexes and 2:1 hINO80-nucleosome complexes, respectively. No peak corresponding to a free INO80 dimer was detected, supporting a model in which two hINO80 complexes associate sequentially on the nucleosome rather than binding as a preassembled dimer.

Together, these results identify two non-exclusive mechanisms that could support bidirectional nucleosome translocation. A single hINO80 complex may reverse direction by switching between alternative nucleosome-binding orientations, while sequential recruitment of a second, oppositely oriented complex could permit directional switching through alternation of the productively engaged motor. Combined with our observation that neighboring nucleosomes confine and reverse translocation, these findings suggest that dynamic reorientation and dual occupancy enable hINO80 to reposition nucleosomes rapidly within defined chromatin boundaries.

## Discussion

INO80 helps organize chromatin across gene bodies and participates in multiple DNA-repair pathways by repositioning nucleosomes to expose damaged DNA and facilitate the recruitment of repair factors^13–15^. However, the mechanism by which INO80 locates and slides nucleosomes within chromatin remains poorly understood. Here, we have combined optical tweezers with single-molecule fluorescence microscopy to investigate hINO80 on naked and chromatinized DNA. On naked DNA, hINO80 undergoes free 1D diffusion, with salt and nucleotide conditions regulating its mobility and DNA-bound lifetime. On chromatinized DNA, INO80 is highly processive, sliding nucleosomes over kilobase-pair distances in rapid, ATP-dependent bursts and reversing direction between successive bursts. Neighbouring nucleosomes act as barriers that confine this movement, causing hINO80 to shuttle a nucleosome repeatedly within a defined region of chromatin. Finally, single-molecule FRET has identified two non-exclusive mechanisms that can support directional reversal: reorientation of a single nucleosome-bound hINO80 complex and alternating engagement of two oppositely oriented complexes bound to the same nucleosome.

The free 1D diffusion of hINO80 provides a potential mechanism for locating nucleosomal substrates after binding within an exposed DNA region. At promoters, hINO80 could bind within the NDR and scan the DNA to locate the adjacent +1 nucleosome. This strategy resembles that of SWR1, which uses ATP-regulated 1D diffusion to locate flanking nucleosomes^41^. Thus, facilitated diffusion may be a conserved target-search mechanism among remodelers^42^.

INO80 occupied at least two diffusive states and could switch between them within a single trajectory. Therefore, the fast and slow states are likely to reflect reversible changes in the interaction between INO80 and DNA. INO80 contains several DNA-binding modules^16,17,30,31,43^, and the slow state may involve more extensive DNA interactions, as increasing the salt concentration shortens the DNA-bound lifetime and favours faster diffusion. Furthermore, diffusion was inversely related to DNA-bound lifetime, supporting a model in which more extensive DNA interactions stabilize binding while constraining movement. Nucleotide state also altered both mobility and lifetime, suggesting that nucleotide-dependent conformational changes regulate the extent of DNA engagement.

On chromatinized DNA, our data show that hINO80-mediated nucleosome sliding is considerably faster and more processive than previously observed. We identified two kinetic populations, with nucleosome sliding speeds of 270 bp/s and 2.3 kbp/s. The slower population agrees well with smFRET measurements showing that INO80 moves nucleosomes by 25 to 50 bp within ∼100 ms, corresponding to rates above 250 bp/s^24,37^. The faster population approaches the velocities of highly processive DNA motors such as Rad54, RecBCD and FtsK, which translocate at 0.5 - 5 kbp/s or faster^44–46^. By comparison, the Snf2-family remodelers RSC and ISW2 can move nucleosomes processively over kilobase distances but at a slower speed of ∼30 bp/s^47^, whilst ACF is less processive (∼200 bp) and much slower (2 bp/s)^48^. The molecular basis of these two hINO80 populations remains unclear. They could represent different INO80 stoichiometries, although differences in substrate composition, such as nucleosome versus hexasome translocation^21^, may also contribute. Previous studies using single nucleosomes positioned on short 601-containing^49^ DNA fragments detected processive sliding over only tens of base pairs and showed strong regulation by flanking DNA length^16,24,37^. By assembling nucleosomes stochastically on λ-DNA, our assay provides kilobases of flanking DNA without a nearby free end and therefore reveals a previously inaccessible regime of hINO80 activity.

Nucleosome sliding occurred in bursts separated by pauses, suggesting that rapid movement is interrupted by transitions into a temporarily inactive or uncoupled state. Static nucleosomes formed robust boundaries that the translocating hINO80-nucleosome complex did not bypass. In contrast, on the intervening naked DNA, the complex could either cross recurrent sites or reverse direction. These intermediate sites may reflect local DNA sequence or variations in DNA mechanics, both of which have been implicated in regulating INO80 activity^46,47^. Despite their different physical origins, pauses at nucleosomal boundaries and intermediate DNA sites exhibited the same kinetics, suggesting that the obstacle determines which direction remains accessible but not the rate at which movement resumes. Departure from either pause may instead require an intrinsic transition within the hINO80-nucleosome complex, such as recoupling of ATP hydrolysis to translocation, reorientation of the complex or switching between active motors. At an intermediate site, both directions remain available, and movement resumes without significant directional bias. At a nucleosomal boundary, continued forward movement is excluded, directing the complex away from the obstacle.

Previous nucleosome-spacing models provide a framework for interpreting the interaction between long-range translocation and chromatin boundaries. The ATP-utilizing chromatin assembly and remodeling factor (ACF) has been proposed to compare the DNA lengths on opposite sides of a nucleosome and bias movement towards the longer flank through coordinated activity of two remodeler complexes^52,53^. hINO80 has also been proposed to act cooperatively, through a mechanism involving two INO80 complexes that monitor both DNA flanks^16^.

Our data suggest two possible mechanisms for bidirectional sliding. First, the smFRET data are consistent with a single hINO80 complex switching between opposite sides of a nucleosome, as proposed for other chromatin remodelers^26,38,39^. Reorientation during a pause would position the motor to resume activity in the opposite direction. However, additional measurements will be needed to demonstrate this. Second, fluorescence intensity analysis and mass photometry showed that two hINO80 complexes can occupy the same nucleosome concurrently. Importantly, the absence of a detectable free hINO80 dimer in solution supports that the 2:1 complex assembles by sequential recruitment rather than binding of a preassembled dimer. This is consistent with the functional 2:1 stoichiometry inferred biochemically^16^. If the two hINO80 complexes adopt opposing orientations, switching which motor is productively coupled to translocation would reverse the sliding direction.

These mechanisms need not be mutually exclusive. Reorientation of a single hINO80 complex would facilitate reversal when hINO80 occupancy is low, whereas dual occupancy could allow rapid motor switching when hINO80 is abundant. The latter may be particularly advantageous at chromatin boundaries, where a poised, oppositely oriented motor could drive movement immediately after a collision.

The combination of diffusion, rapid translocation and barrier sensing may be relevant to several chromatin contexts. At promoters, hINO80 could scan an NDR, locate the +1 nucleosome and reposition it until movement is constrained by neighbouring nucleosomes or other chromatin-bound factors. Local flanking-DNA sensing could then define its final position and help propagate nucleosome spacing into the gene body. Long-range translocation may be particularly important at sites of DNA repair, where nucleosomes must be removed from regions considerably larger than typical promoter NDRs. Moving nucleosomes over kilobase-pair distances would allow rapid access for damage recognition, end processing, and repair-complex assembly. In cells, transcription factors, polymerases, cohesin and other DNA-bound complexes may also act as barriers. Together, our findings redefine hINO80 as a dynamic, reversible motor whose powerful long-range nucleosome sliding activity is shaped by the surrounding chromatin to achieve controlled nucleosome positioning.

## Acknowledgements

We thank Prof. Dale Wigley and Dr Adam S.B. Jalal for providing materials, critical reading of the manuscript and valuable discussions regarding the broader context of our findings. Funding: the work was funded by a core grant from the MRC Laboratory of Medical Sciences (UKRI MC-A658-5TY10 [D.S.R.]).

## Author contributions

B.A., P.G., and D.S.R. designed the studies. B.A. performed and analyzed the single-molecule experiments, and P.G. and M.T.S. conducted and analyzed the biochemical experiments. P.G., M.T.S., E.A.M. and D.I.E. prepared the samples. B.A., P.G., and D.S.R. analyzed the data and wrote the manuscript with input from all the authors.

## Methods

### Purification of fluorescently labelled hINO80

hINO80 complex was purified as previously described^19^. To label hINO80 specifically on the Ies6 subunit, an hINO80 complex lacking the ARP5/IES6 module (hINO80ΔARP5IES6) was cloned, expressed and purified similar to the complete complex. Genes encoding human ARP5 with a twin StrepII tag and IES6 with a ybbR tag were cloned into a multibac expression vector and expressed in insect cells. The ARP5-2xSII and IES6-ybbR module was purified as follows, cells were resuspended in Buffer A (25 mM HEPES, pH 7.5, 300 mM NaCl, 1 mM TCEP, 10% glycerol) supplemented with protease inhibitor tablets and benzonase and lysed by sonication. After clarifying the lysate by centrifugation followed by filtration, the supernatant was injected onto a StrepTrap HP column, washed with Buffer A, and eluted in Buffer A supplemented with 2.5 mM desthiobiotin. The eluted protein was combined and diluted with 25 mM HEPES pH 7.5, 1 mM TCEP, 10% glycerol to reduce the NaCl concentration to 150 mM. The sample was loaded onto a HiTrap Q HP (Cytiva) column, washed with Buffer B (25 mM HEPES pH 7.5, 150 mM NaCl, 1 mM TCEP, 10% glycerol) and eluted with a gradient of 5% to 100% Buffer C (25 mM HEPES pH 7.5, 2 M NaCl, 1 mM TCEP, 10% glycerol) over 10 mL. The ARP5/IES6 module was labelled using Atto 647N-CoA and Sfp and then purified using a HiTrap Heparin HP (Cytiva) column with the same buffers as used for the HiTrap Q HP purification step. Labelled ARP5/IES6 was combined with INO80ΔARP5IES6 to reconstitute the conserved core INO80 complex, which was then further purified using a HiTrap Heparin HP (Cytiva) column with the same buffers as used for the HiTrap Q HP.

### Preparation of histone octamer

Human histones, H2A (or H2A(K118C) for fluorescent labelling), H2B, H3.1 and H4 were expressed and purified as previously described, with minor modifications^19^. Briefly, H2A and H2B dimers were co-expressed in E. coli, cells were lysed by sonication in lysis buffer (20 mM Tris pH7.5, 500 mM NaCl, 0.1 mM EDTA, 1 mM TCEP, supplemented with Roche Protease Inhibitor tablet). Dimers were purified as soluble proteins using HiTrap Q FF and HiTrap Heparin HP columns in buffer A (20 mM Tris pH 7.5, 500 mM NaCl, 1 mM EDTA, 1 mM TCEP) and eluted, after removal of the HiTrap Q FF columns, using a gradient to 100% buffer B (20 mM Tris pH 7.5, 2 M NaCl, 1 mM EDTA, 1 mM TCEP), followed by gel filtration on Superdex 200 using buffer A. For labelled dimers, H2A(K118C) H2B dimers were labelled with Alexa Fluor 555 and then repurified using a 1ml HiTrap Heparin column (100%B step elution). Tetramers of H3.1 and H4 were co-expressed in E. coli and lysed in lysis buffer. Tetramers were purified as soluble proteins on HiTrap Heparin HP in buffer A, eluting using a gradient to 100% buffer B, followed by gel filtration on Superdex 200 using buffer B. Histone octamers were prepared by mixing tetramers with a 10% molar excess of dimers and then purified by gel filtration on Superdex S200.

### Nucleosome preparation

For the TIRF and mass photometry experiments, a biotin-113-widom601-2 nucleosome with an Alexa Fluor 555 at SHL2.5 was prepared as follows. An AvaI-StyI fragment was prepared by annealing and ligating together oligo pairs 853/863 and 855/929 followed by purification on 8 ml MonoQ column using a 20-30%B gradient, where Buffer A is 20 mM Tris 7.5, 1 mM EDTA, 250 mM NaCl, and Buffer B is 20 mM Tris 7.5, 1 mM EDTA, 2 M NaCl. A 59mer with C6-Amino modification was labelled using Alexa Fluor 555 N-hydroxysuccinimide ester and purified by HPLC. This 59mer was annealed to a 57mer to create a StyI-AccI fragment, which was ligated to the AvaI-StyI fragment. Purification on 8 ml MonoQ column (25-32%B) separated labelled AvaI-AccI fragment from unlabelled. Oligo pair 710/1065 (AccI-RV fragment) was annealed and ligated to labelled AvaI-AccI fragment and purified on 8 ml MonoQ column (25-32%B).

Nucleosomes were reconstituted from octamers (described above) and the AvaI-RV fragment by salt gradient dialysis in several steps from 2 M to 0.2 M NaCl. A 101 bp biotinylated fragment was prepared by PCR (with Biotin-primer and Primer-2) on a plasmid containing 24 copies of a target sequence as previously described^19,54^ and digested with AvaI. This 101 bp fragment was purified by ion-exchange chromatography on a HiTrap Q FF column, and ligated to the above nucleosome and finally purified by glycerol gradient ultracentrifugation. Oligos referenced here are described in the table below

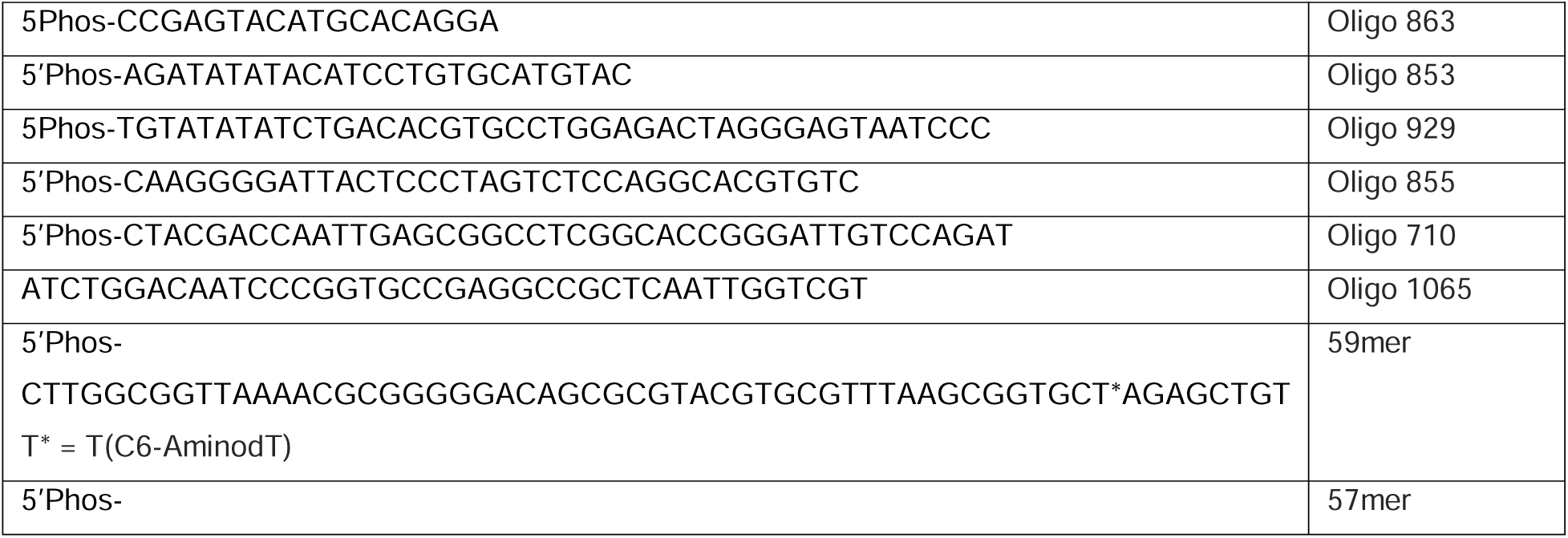
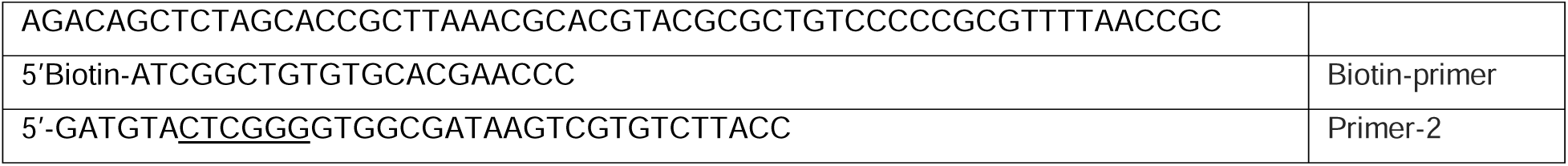
Table of Oligos.

### yNAP1 preparation

*S. cerevisiae* yNAP1 containing an N-terminal His tag and TEV cleavage site in a pRSETa vector was expressed and purified from *E. coli*. Expression was induced by the addition of 0.5 mM IPTG and harvested after 4 h at 37°C. Cells were lysed in buffer A (50 mM HEPES pH 8.0, 150 mM NaCl, 1 mM DTT) supplemented with EDTA-free protease inhibitor tablet (Roche) using sonication. The clarified lysate was purified on a HisTrap HP column in buffer A using a step elution to buffer B (50 mM HEPES pH 8.0, 150 mM NaCl, 1 mM DTT, 500 mM imidazole). Peak fractions were pooled and further purified on a HiTrap Q FF column in buffer C (50 mM HEPES, pH 8.0, 100 mM NaCl, 1 mM DTT) and eluted using a gradient to buffer D (50 mM HEPES, pH 8.0, 2 M NaCl, 1 mM DTT). Fractions were pooled, concentrated and further purified using a Superdex 200 Increase 10/300 GL gel filtration column in buffer E (50 mM HEPES, pH 8.0, 500 mM NaCl, 1 mM TCEP). Fractions were pooled, concentrated, and stored at −80°C.

### Correlative optical tweezers with fluorescence detection

Single-molecule optical tweezer experiments were carried out using a commercially available C-trap system (Lumicks). Before use, the microfluidic system was passivated with BSA (0.2 mg/ml) and Pluronics F-127 (0.5% w/v) in 50 mM Tris-HCl, 7.5 and 50 mM NaCl, flowing at least 500 μl through all channels. SPHERO Streptavidin Coated polystyrene beads (4.35 μm) at 0.005% w/v were flowed in channel 1, caught using twin optical traps with a target stiffness of 0.28 pN/nm. Biotinylated λ-DNA, flowed in channel 2, was caught between the traps and checked against a worm-like chain model. For experimental acquisition, the beads DNA, and any bound proteins were brought to channel 3, containing experimental buffer of 25 mM Tris-HCl, pH 7.8, 100 mM KCl, 4% Glycerol, 2 mM MgCl_2_, 1 mM EDTA.

For experiments with hINO80 on naked DNA, Atto 647N labelled hINO80 (1 nM) was flowed into channel 4. Caught λ-DNA was held at 10 pN and taken to channel 4, where it was incubated with a live kymograph running (638 nm laser, 35 μW, 100 nm pixel size, 0.1 ms per pixel, ∼33 ms line time) to check for binding of INO80 molecules. Once ∼3 hINO80 molecules had bound, we moved the tether to channel 3 to continue imaging the kymograph and monitor diffusion until all INO80 molecules had photobleached. Nucleotides ATP and ADP were added at 1 mM concentration. The non-hydrolyzable analog, ADP.BeFx, used 3 mM ADP, 3 mM BeCl_2_, and 15 mM NaF.

For the nucleosome force-spectroscopy experiments, a human histone octamer with AF555 labelled H2A (∼1 nM), and yNap1 (1 nM), were flowed into channel 4. Caught λ-DNA was held at ∼1 pN and taken to channel 4, and incubated for 1-5 s. DNA was then returned to channel 2 and briefly washed under low flow to remove any free unwrapped histones and yNap1 from the λ-DNA tether. Nucleosome density on the DNA was checked via 1D confocal imaging with a 532 nm laser (5 μW), and incubated for 1-5 s to obtain 3-5 nucleosomes per tether. The partially chromatinised DNA was then taken to channel 3, and pulled at 50 nm/s to acquire force-extension curves.

For Nucleosome photobleaching experiments, the above was repeated with confocal imaging at higher laser power (12.5 μW) to produce kymographs until all tethered nucleosomes had photobleached.

For experiments with both hINO80 and nucleosomes, the nucleosomes were prepared as above, at 1 pN, with the octamer and yNap1 in channel 4, then the partially chromatinised DNA was incubated with hINO80 in channel 5, before being taken back to channel 3 for recording (532 nm laser μW, 638 nm laser 35 μW). Channel 3 contained either the experimental buffer as described above (no nucleotide), with 1 mM ATP, or with 1 mM ATP and 0.1 mM ADP. Kymographs were taken with a line time of 33 ms, whereas 2D videos were taken at a 1-second frame time.

In all cases, force-extension curves and kymographs were exported as HDF5 files from bluelake for analysis.

### Optical tweezers analysis

Optical tweezers data were analyzed using custom Jupyter notebooks built using the Pylake-Python package. For force spectroscopy, the low-frequency force and distance data were extracted, and a peak-finding algorithm was used to identify ruptures in the curve, indicative of nucleosome unwrapping. These were used to categorize the pre- and post-unwrapping force-extension curves, which were then fit with a piecewise extensible worm-like chain model (eWLC):

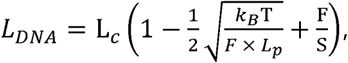

where the persistence length (L_p_), and stretching modulus (S) are constant between pieces, but each piece has a distinct contour length (L_C_). The contour length increase between pieces (ΔL_C_) corresponds to the DNA length released by nucleosome unwrapping.

To track diffusing hINO80 molecules, kymographs were extracted and downsampled by 3-fold (final frame time ∼100 ms) to correct for blinking molecules in the trace. Tracks were initialised by hand-drawn tracing, omitting collisions with other molecules or the bead. To localize the molecule’s position, each frame of the downsampled initialised track was fit independently with a Gaussian point spread function^55^ using the refine_tracks_gaussian method in Pylake with a refinement window of 5 pixels, yielding a fitted trajectory. To obtain a trajectory of diffusion coefficients against time, we employed a rolling window method similarly to previously described^28,29^. A rolling window over the position trajectory is used to compute mean squared displacement (MSD) for increasing time separations (tau) with a maximum tau of 10 frames, within a downsampled 50-frame window, and MSD against tau is fit with a straight line (with offset, for localization error). Diffusion coefficients were then obtained from the gradient using the relation

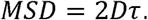

Any gradients with a negative gradient were discarded, rather than converted to negative diffusion coefficients. Because this method produces a diffusion coefficient trajectory shorter than the position trajectory, it is plotted with a 25-frame offset. Diffusion coefficients obtained this way are converted from μm^2^/s to bp^2^/s using the known length of *λ*-phage genome (48,502 bp) and the physical tether length between the centers of beads in the kymograph, subtracting the bead diameter.

Diffusion coefficients collected from each trajectory were plotted in histograms for each condition in log space, then fit with log Gaussians using scipy.optimize curve_fit. The two Gaussian fits of the 100 mM KCl condition (no ATP) were used to define the cutoff between the slow and fast states as the midpoint (in log space) between the two Gaussian centers. The cutoff between static and slow distributions in the ADPBeF condition was defined similarly. To discern free diffusion from processivity in the Apo and ATP conditions, MSD analysis was performed on whole traces with a freely fit anomalous diffusion factor as follows

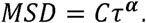

For nucleosome photobleaching trajectories, static tracks were localized as above, extending each track beyond the photobleaching point to obtain a baseline. Photon intensities were extracted and classified into single- or double-step photobleaching.

For hINO80-induced nucleosome sliding, kymographs were analyzed raw (not down-sampled) due to the rapid motion of the complex. hINO80 tracks (red channel) and nucleosome tracks (green channel) were localized as described above. For the purposes of colocalization, bouncing, and crossing, a threshold boundary of 250 nm on either side of the mean position of a static nucleosome was used to assign colocalization.

To quantify nucleosome sliding bursts, sections of trajectories were identified in the ATP and ATP+ADP conditions where the burst was flanked by a pause before and after to help anchor a static part of the trajectory, and intensities were extracted from the Atto 647 (hINO80) channel. These were fit with a piecewise function (scipy.optimize curve_fit) comprised of a 1D Gaussian point spread function with fixed amplitude and width, that moves at a constant velocity from p1 at t1 to p2 at t2 (**Fig. 2G**). The Gaussian has across all three parts of the trajectory. The parameters p1, t1, p2, and t2 are initialized by hand via clicking on the kymograph image, but then allowed to freely fit to the intensities of each pixel, along with the global gaussian width and amplitude.

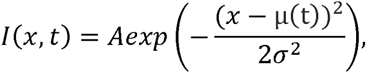

where

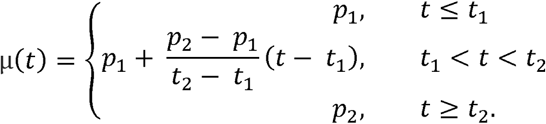

Burst durations were plotted against burst distance and fit with a two-component mixture of linear regressions where each population has its own independent slope and intercept, where cluster membership and regression parameters were obtained jointly by expectation-maximisation. Fit was repeated with 60 random initialisations and the solution with highest log-likelihood retained. Standard errors for the slope were obtained from asymptotic covariance of weighted least-squares fit.

Pausing analysis was done by extracting trajectories from the hINO80 channel on 3 tracks that exhibited clear repeated pausing of bursts between barrier nucleosomes. k-means clustering on the position trajectory identified the two edge positions, as well as intermediate pausing positions in one trajectory. Dwell-time analysis used exponential fitting to dwells longer than one frame (omitted as either noise or the molecule passing through a state without stopping). Events at intermediate pausing positions were classified as “continue” if the molecule arrived from one direction and left in the other, and conversely classified as “reverse” if it returned in the direction it came.

In all cases in this paper, fluorescence images show the “red” channel (Atto 647N, hINO80) in orange, and the “green” channel (AF 555, H2A) in blue, with coincidence of the two appearing white.

### TIRF data acquisition

Total Internal Reflection Fluorescence (TIRF) data were acquired with a home-built microscope, as described^26^. Briefly, fluorophores were excited with a 532 nm laser (Stradus, Vortran) and a 637 nm laser (Stradus, Vortran) in alternating laser excitation (ALEX) mode. Fluorescence was collected through a 1.2 NA, 60x water objective (Olympus) and filtered through a dual bandpass filter (FF01-577/690-25, Semrock). Donor and acceptor fluorescence was separated by spectrally filtering using an OptoSplit II (Cairn Research), and further filtered through ET585/65M and ET700/75M (Chroma) bandpass filters. Donor and acceptor images were projected side-by-side onto an EMCCD (Andor iXon Ultra 897). Data were collected as raw movies using a custom LabVIEW script at 100 ms time resolution.

### Flow chamber assembly and assay

Quartz slides (UQC optics) and glass coverslips were aminosilinized with N-(2-Aminoethyl)-3-aminopropyltrimethoxysilane, then pegylated using methoxy-PEG-SVA (Mr = 5,000, Laysan Bio, Inc.) containing 5 % biotin-PEG-SVA (Mr = 5,000, Laysan Bio, Inc.) in 100 mM sodium bicarbonate as described with minor modifications^26^. Following passivation, slides and coverslips were stored under nitrogen in the dark at −20°C. Prior to use, slides and coverslips were warmed to room temperature and assembled into flow chambers using 0.12 mm thick double-sided adhesive sheets (Grace Bio-Labs SecureSeal). Flow chambers were sealed with epoxy glue.

Biotinylated 113N2 nucleosomes with AF555 at SHL2.5 (described above) were injected into the assembled flow chamber, and excitation with 532 laser was used to assess density. Atto 647N labelled INO80 (5 nM) was injected in imaging buffer (100 mM Tris-HCl, pH 7.8, 100 mM KCl, 4% glycerol, 2 mM MgCl_2_, 1 mM EDTA, 0.2 mg/ml BSA, 2.5 mM protocatechuic acid, 0.25 μM protocatechuate-3,4-dioxygenase, 1 mM 4-nitrobenzyl alcohol, 1 mM ascorbic acid, 2 mM methyl viologen). Data were then acquired at 100 ms frame time, alternating the 532 nm and 638 nm lasers between frames.

### TIRF Data Analysis

Single-molecule fluorescence spots from the raw movies were localised using custom IDL scripts and converted into raw fluorescence trajectories^56^. Raw fluorescence trajectories were corrected for bleed-through of the donor fluorescence into the acceptor channel (alpha correction factor). Apparent FRET efficiencies were calculated as the ratio of acceptor intensity divided by the sum of the donor and acceptor intensities under donor excitation. Emission of the acceptor under excitation was also extracted.

Extracted traces were analyzed using a custom MATLAB program to select trajectories with hINO80 binding. Trajectories were further analysed with custom Jupyter notebook Python scripts. Emission intensities under 532 nm (donor) and a 637 nm (acceptor) excitation were extracted from raw traces and despiked using a Hampel filter (rolling median, 5-MAD threshold, 11-frame window). Acceptor excitation traces were normalized per-molecule to a common intensity scale using a robust baseline (1st percentile) and plateau reference (maximum of a 15-frame smoothed trace). FRET traces were calculated as I_A_/(I_D_ + I_A_) and background-corrected by identifying each trace’s photobleaching point (single-change-point detection on donor intensity) and subtracting the post-bleach baseline per channel; traces were truncated at the bleach point. States were assigned by fitting a consensus variational-Bayes hidden Markov model across all traces of each condition simultaneously^57^. Model order was selected per dataset by comparing fits with different numbers of states; redundant background states were merged by summing the corresponding blocks of the transition-count matrix. Dwell times were obtained from the Viterbi-decoded state path. Survival functions were fit with a double-exponential model; transitions confounded by a competing outcome (e.g., dimerization pre-empting monomer dissociation) were treated as right-censored (Kaplan-Meier) rather than as departure events. Rate uncertainties were estimated by bootstrapping over individual dwell events (2000 resamples) and reported as robust (MAD-based) standard errors.

### Mass photometry

Mass photometry experiments were carried out using a commercial system (Refeyn Two MP). Mass calibration was carried out using thyroglobulin. Sample carrier slides (Refeyn) were used with gaskets (Grace-Bio-Labs, CultureWell 50 - 3 mm). Atto 647N labelled hINO80 and the 113N2 nucleosomes were diluted to 40 nM and 20 nM, respectively, in experimental buffer (25 mM Tris-HCl, pH 7.8, 100 mM KCl, 4% glycerol, 2 mM MgCl_2_, 1 mM EDTA), and incubated for 2 minutes to allow complexes to form. Concentrated solution was diluted 1:20 into a buffer droplet on the instrument, and then data were acquired using large field of view settings (AcquireMP) for 300 s. DiscoverMP software was then used to apply the mass calibration to the data and masses were exported. Mass data were then fitted with Gaussians using custom Jupyter notebook Python scripts with scipy.optimize curve_fit.

