## Supplementary Materials for "INO80 rapidly shuttles nucleosomes between chromatin barriers"

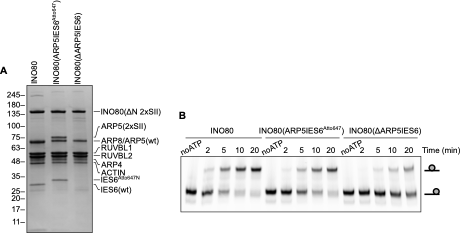

**Supplementary Figure 1. Preparation of labelled hINO80.** **A.** SDS-PAGE of hINO80 showing each subunit (left), reconstituted with labelled ARP5/IES6 (middle) and with ARP5/IES6 deleted (right). **B**. Electrophoretic mobility gel-shift assay of hINO80 sliding a nucleosome. Labelled hINO80 has comparable activity to unlabeled. Deletion of ARP5/IES6 impairs remodelling activity.

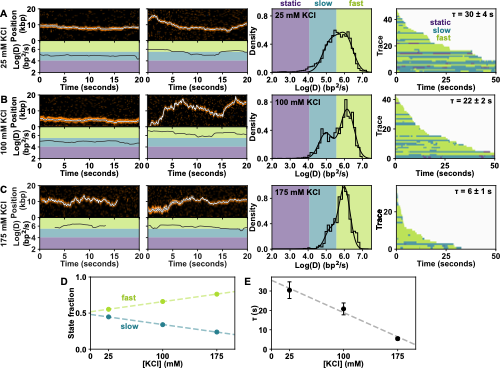

**Supplementary Figure 2. Salt titration of hINO80 diffusion on naked DNA.** Example kymographs (top) and rolling-window diffusion coefficients (bottom) of hINO80 on naked DNA in (**A**) low-salt (25 mM KCl), (**B**) standard condition (100 mM KCl) and (**C**) high-salt (175 mM KCl). Distribution of rolling-window diffusion coefficients (third column). Shaded regions indicate static (purple), slow (teal), and fast (green) regimes (as in **Fig. 1F**). Rastergram of all trajectories (fourth column) ordered by length, colored by diffusion category as above, with corresponding dissociation lifetimes (with s.e.m.). **D.** Fraction of trajectory in either the fast or slow regimes (from Gaussian amplitudes) as a function of salt concentration. **E.** Dissociation lifetimes (from rastergrams) as a function of salt concentration.

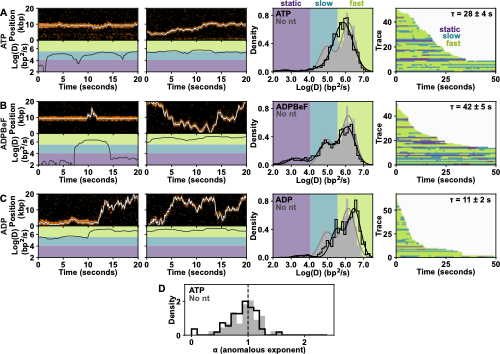

**Supplementary Figure 3. Effect of nucleotides on hINO80 diffusion on naked DNA.** Example kymographs (top) and rolling-window diffusion coefficients (bottom) of hINO80 on naked DNA in (**A**) 1 mM ATP, (**B**) ADPBeF and (**C**) 1 mM ADP. Distribution of rolling-window diffusion coefficients (third column). Shaded regions indicate static (purple), slow (teal), and fast (green) regimes (as in **Fig. 1F**). Rastergram of all trajectories (fourth column) ordered by length, colored by diffusion category as above, with corresponding dissociation lifetimes (with s.e.m.). **D.** Distribution of anomalous exponents from the mean-squared-displacement analysis for all trajectories in the presence (white) and absence (grey) of ATP, confirming unconstrained 1D Brownian motion.

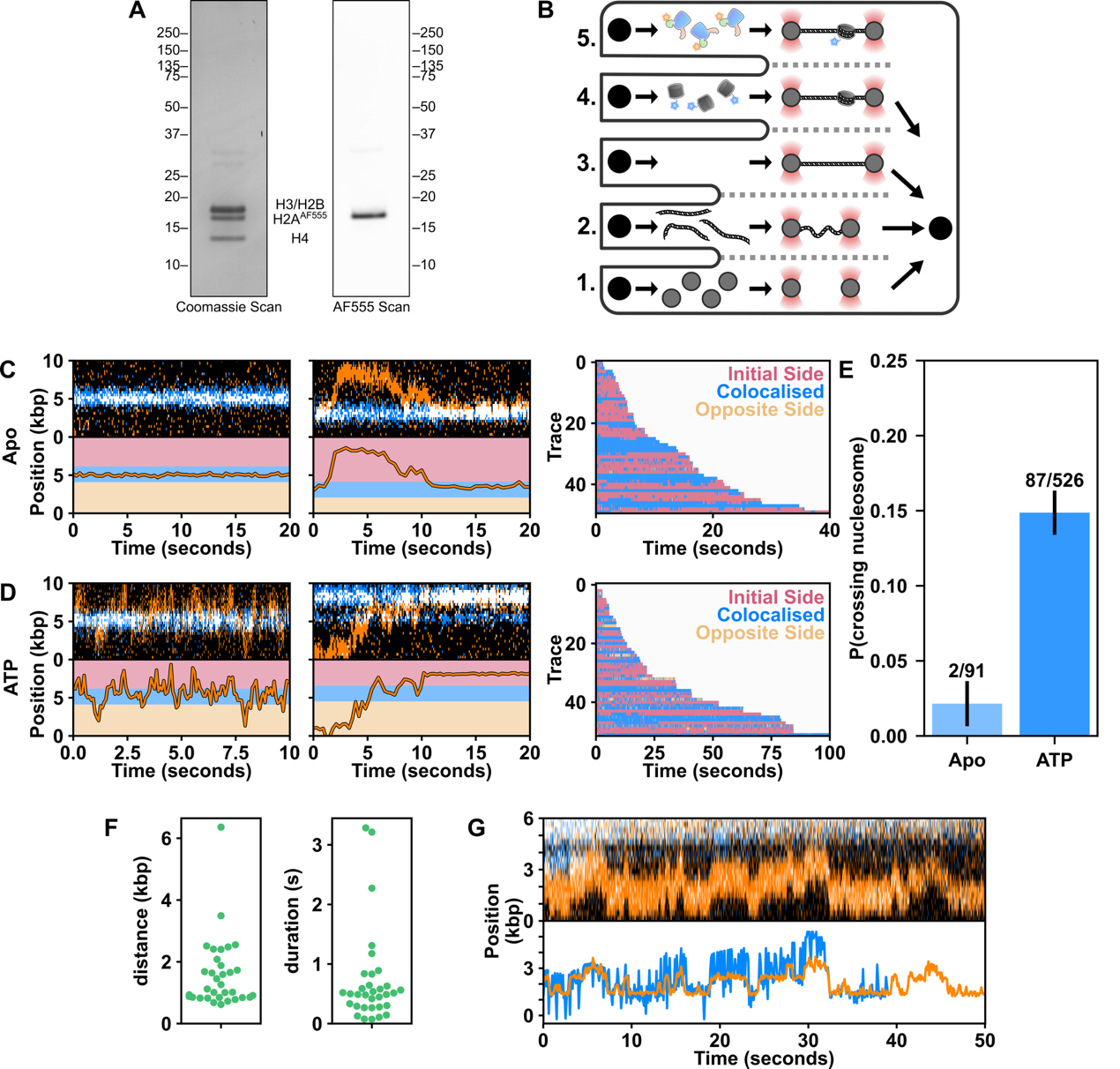

**Supplementary Figure 4. Dynamics of hINO80 on chromatinized DNA.** **A.** SDS-PAGE of labelled human histone octamer (left: Coomassie-stained, right: AF555 fluorescence). **B.** Optical tweezers laminar flow cell with streptavidin beads (Channel 1), biotinylated λ-DNA (Channel 2), imaging buffer (Channel 3), labeled human histone octamer (Channel 4), and labelled hINO80 (Channel 5). **C.** Example kymographs (top) and hINO80 localization (bottom) in the absence of ATP, showing colocalization with a nucleosome (left) and free diffusion and binding to a nucleosome (right) but no sliding. Regions beyond 250 nm on either side of the nucleosome are shaded pink or yellow (bottom) to highlight the absence of nucleosome crossing. Rastergram of all trajectories colored by position of hINO80 relative to nucleosome (third column), highlighting very few crossings (likely due to localization error). **D.** Example kymographs (top) and hINO80 localization (bottom) and rastergram in the presence of ATP (similar to **C**), showing nucleosome crossing. **E**. Nucleosome crossing probabilities in the presence and absence of ATP (binomial standard errors). **F** Distributions of size (distance), and duration of hINO80 nucleosome sliding bursts. **G.** Additional example kymograph (top) and localization (bottom) of hINO80 (orange) repeatedly sliding a nucleosome back and forth between two flanking nucleosomes (blue) as in Figure 3A of the main text.

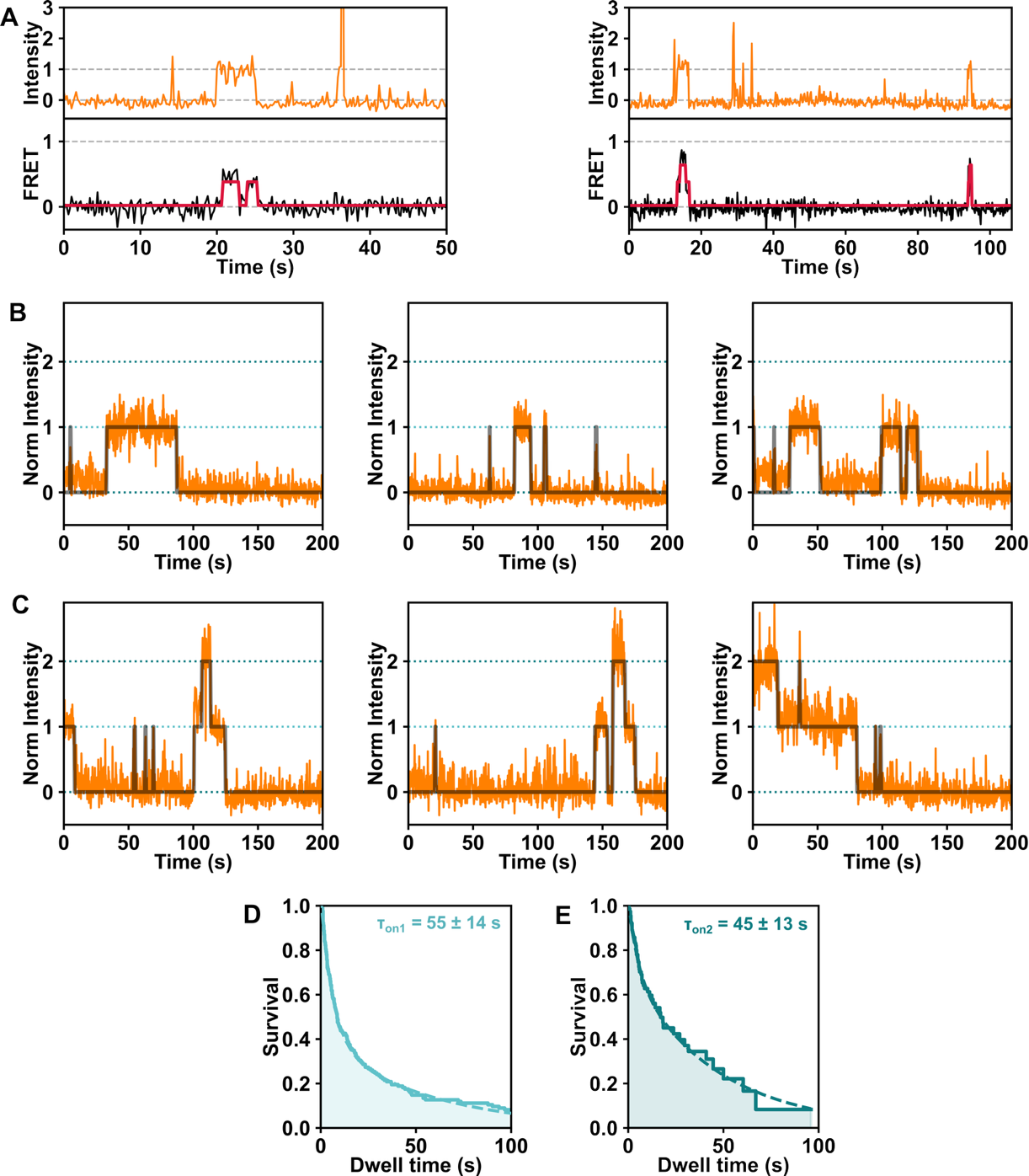

**Supplementary Figure 5. Example FRET and fluorescence intensity trajectories.** **A.** Single-molecule trajectories showing initial hINO80 event (top) and subsequent FRET dynamics (bottom). **B.** Single-molecule fluorescence trajectories of labeled hINO80 binding a surface-immobilized mono-nucleosome, showing 1:1 complex formation. **C.** Single-molecule fluorescence trajectories of labeled hINO80 binding a surface-immobilized mono-nucleosome, showing 2:1 complex formation. Dwell time distributions in the unbound state (**D**) and before binding of a second hINO80 (**E**), yielding the binding times of the first and second hINO80, respectively. The two times are within error of each other, indicating sequential recruitment rather than cooperative binding.
